# Entropy in Primary Sensory Areas Lower than in Associative Ones: The Brain Lies in Higher Dimensions than the Environment

**DOI:** 10.1101/071977

**Authors:** James F. Peters, Arturo Tozzi, Ebubekir İnan, Sheela Ramanna

## Abstract

The brain, rather than integrate sensory inputs and concentrate them into concepts as currently believed, appears to increase the complexity from the perceived object to the idea of it. Topological models predict indeed an increase in dimensions and symmetries from the environment to the higher activities of the brain. Models predict that informational entropy in the primary sensory areas must be lower than in the higher associative ones. In order to demonstrate the novel hypothesis, we introduce a method for the measurement of information in fMRI neuroimages, *i.e*., nucleus clustering’s Rényi entropy derived from strong proximities in feature-based Voronoï tessellations, *e.g.,* maximal nucleus clustering. The technique facilitates the objective detection of entropy/information in zones of fMRI images generally not taken into account. We found that the Rényi entropy is higher in associative cortices than in the visual primary ones. It suggests that the brain lies in higher dimensions than the environment and that it does not concentrate, but rather dilutes the message coming from external inputs.

---

Sequential processing of information is hierarchical, such that the initial, low-level inputs of primary sensory areas are transformed into representations and integration emerges at multiple processing associative cortical stages (Werner and Noppeney 2009; Nieuwenhuys et al. 2008). Brain functions organize in global gradients of abstraction and a spatially progressive increases in amalgamation of representation or function emerge as cortical distance from the input increases (Taylor et al., 2016). In sum, the current paradigm talks about a brain that, through the limited human sensory repertoire, extracts and concentrates information from the environment (**Figure 1A**). However, recent topological advances talk about a different scenario. The Borsuk-Ulam theorem (BUT) states that (Borsuk 1933):

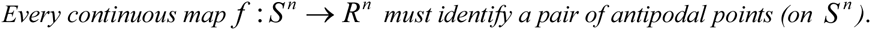

This means that the positive-curvature sphere *S*^*n*^ maps to the Euclidean space *R*^*n*^, which stands for an *n*-dimensional Euclidean space (Matoušek, 2003). Many BUT variants have been recently proposed (Peters and Tozzi, 2016; Tozzi and Peters, 2016a; Tozzi and Peters, 2016b; Peters et al., 2016). A generalized version of BUT states that:

*Multiple sets of objects with matching descriptions in a d-dimensional manifold M^d^ are mapped to a single set of objects in Md-1 and vice versa.* The sets of objects, which can be physical or biological features, do not need to be antipodal and their mappings need not be continuous. The term *matching description* means the sets of objects display common feature values.*M* might stand for manifolds with any kind of curvature, either concave, convex or flat. *M^d-1^* may also be a part of *M^d^*. The projection from *S*^*n*^ to *R*^*n*^ in not required anymore. Instead, just *M* is required. The notation *d* stands for a natural, rational, or irrational number. This means also that the need for natural number and spatial dimensions of the classical BUT is no longer required. There exists a physical link between the abstract concept of BUT and the real energetic features of systems formed by two manifolds *M*^*d*^ and *M^d-1^*. *M*^*d*^ is equipped with two matching points, standing for symmetries. When the two points map to a *n*-Euclidean manifold where *M^d-1^* lies, a symmetry break and dimensionality reduction occurs, and a single point is achieved in the image space. It is widely recognized that a decrease in symmetry goes together with a reduction in entropy and free-energy (in a closed system) (Roldán et al., 2014). This means that the single mapping function on *M^n-1^* displays energy parameters lower than the sum of the two corresponding matching functions on *M^d^*. Therefore, a decrease in dimensions gives rise to a decrease of energy and energy requirements, and vice versa. This *energy-BUT* variant concerns not just energy, but also information. Indeed, two antipodal points contain more information than their single projection in a lower dimension. Dropping down a dimension means each point in the lower dimensional space is simpler, because each point has one less coordinate. In sum, energy-BUT provides a way to evaluate the decrease of energy in topological, other than thermodynamical, terms.

Given the recent claims of brain higher dimensionality (Tozzi and Peters, 2016a), we propose a novel framework where the environment displays less dimensions and symmetries than the central nervous system, so that external inputs progressively increase their complexity when spikes cross the brain towards higher associative cortices (**Figure 1B**). In order to demonstrate our hypothesis, we evaluated, through a novel neuroimaging technique, the entropy values in different cortical areas after visual stimuli presentation.

**Figure.**
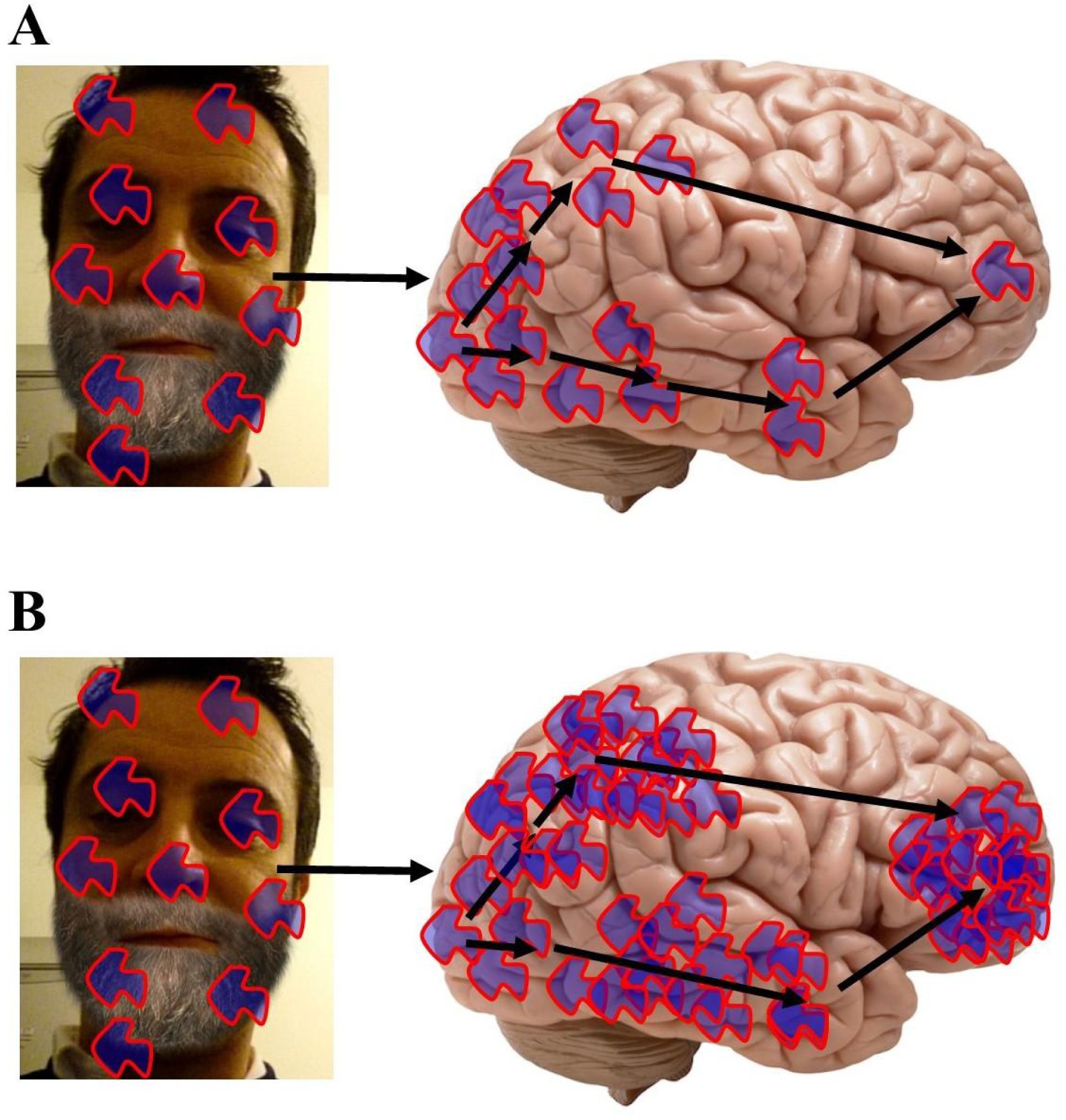
Two models of information processing in the brain. The current paradigm (**Figure 1A**) states that the sensory input is progressively concentrated by areas far from the primary sensory ones. In the higher associative areas, a general concept related to the visual percept is achieved. Therefore, starting from the overwhelming complexity of the environment, the brain focuses just on what it is useful for the individual preservation. The alternative scenario hypothesizes that the sensory input is progressively diluted when passing from the lower sensory areas to the higher associative ones (**Figure 1B**). In other words, the novel hypothesis states that the concept is more complex than the inputs from the external world. The brain activity leads to increases, and not decreases, of complexity, compared with the surrounding environment.

## MATERIALS AND METHODS

### 1 Samples

We evaluated 16 fMRI images from visual tasks experiments, which illustrate the activity of different brain areas during vision. The images, taken from Mandelkow et al. (2016), display a widely distributed fMRI response throughout the brain, elicited by basic vision and object recognition (**Figure 2A**). A voxel-wise ANOVA F-statistic map (threshold p < 1%, uncorrected) was superimposed on the T1-weighted anatomical MRI of one representative subject (16 axial slices in radiological convention). Each tessellated image leads to the MNC mesh clustering described in the next paragraph.

**Figure 2.**
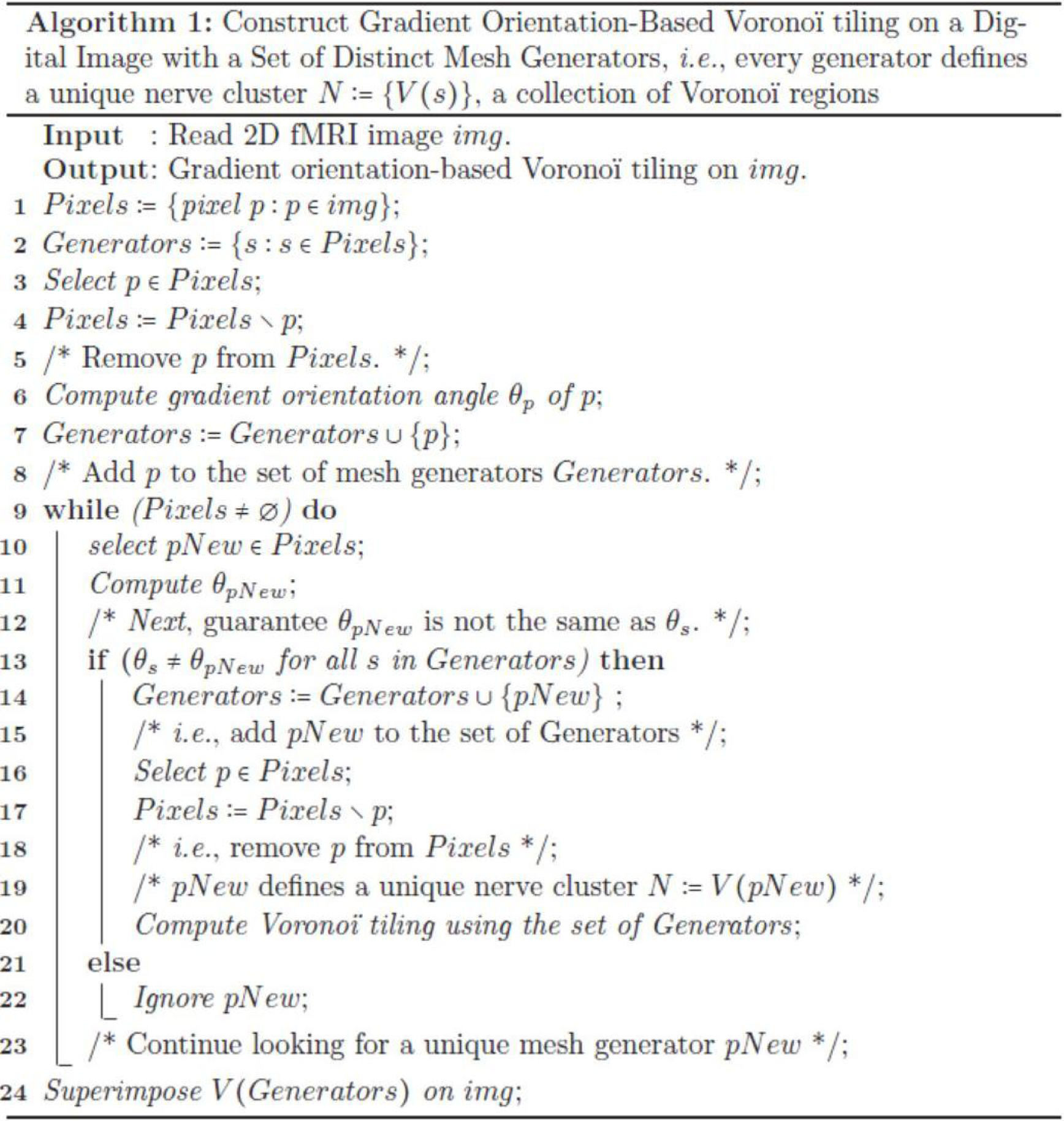
The steps in the method used to construct the mesh on an fMRI image shown in **Figure 1A**.

### 2 Generating Points in Voronoï Tilings of Plane Surfaces

This section introduces nucleus clustering in Voronoï tessellations of plane surfaces (Peters 2016; Edelsbrunner, 2006). A Voronoï tessellation is a tiling of a surface with various shaped convex polygons (Duyckaerts and Godefroy, 2000; Frank and Hart, 2010). Let *E* be a plane surface such as the surface of an fMRI image and let *S* be a set of generating points in *E*, *s* ∈ *S*. Each such polygon is called a *Voronoï region V*(*s*) of a

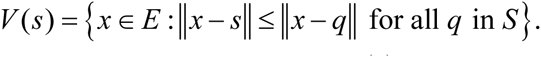

In other words, a Voronoï region *V*(*s*) is the set of all points *x* on the plane surface E that are nearer to the generating point *s* than to any other generating point on the surface (**Figure 2B**). In this investigation of fMRI images, each of the generating points in a particular Voronoï tessellation has a different description. Each description of generating point *s* is defined by the gradient orientation angle of *s*, *i.e*., the angle of the tangent to the point *s*.

### 3 Nucleus Clustering in Voronoï Tilings

A *nucleus cluster* in a Voronoï tiling is a collection of polygons that are adjacent to (share an edge with) a central Voronoï region, called the cluster nucleus. In this work, the focus is on maximal nucleus clusters, which highlight singular regions of fMRI images (**Figure 2C**). A pair of Voronoï regions are considered *strongly near*, provided the regions have an edge in common (Peters and Inan, 2016; Peters, 2016). A *maximal nucleus cluster N* contains a nucleus polygon with the highest number of strongly near (adjacent) Voronoï regions (**Figure 2D**). The gradient orientation angle *θ* of a point (picture element) in an fMRI image is found in the following way. Let *img*(x, y) be a 2D fMRI image. Then

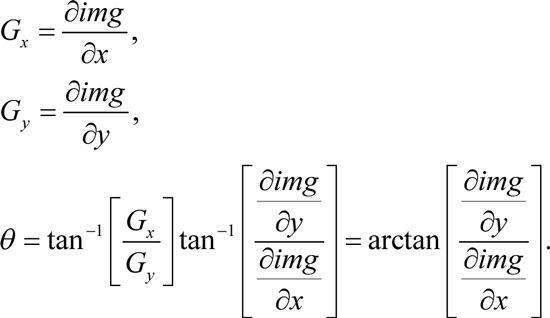

In other words, the angle*θ* of the generating point of the mesh nucleus is the arc tangent of the ratio of the partial derivatives of the image function at a particular point (*x, y*) in an fMRI image.

### 4 Steps to Construct a Gradient-Orientation Mesh

Here we give the steps to build a Voronoï tiling, so that every generating point has gradient orientation (GO) angle *θ* that is different from the GO angles of each of the other points used in constructing the tiling on an fMRI image (see **Figure 2B**). The focus in this form of Voronoï tiling is on guaranteeing that each nucleus of a mesh cluster is derived from a unique generating point. This is accomplished by weeding out all image pixels with non-unique GO angles. The end result is a collection of Voronoï regions that highlight different structures in a tessellated fMRI image. Each Voronoï region *V*(*s*) in a GO mesh is described by feature vector that includes the GO of the generating point *s*.

Since each s is unique (not repeated in the Generators set in **Figure 2B**), each nucleus mesh cluster *N* has a unique description. Taking this a step further, we identify maximal nucleus clusters on a tessellated fMRI image. In effect, each maximal *N* tells us something different about each region of a tiled fMRI image, since we include, in the description of a maximal nucleus, the number adjacent regions as well as the GO of the nucleus generating point.

### 5 Rényi entropy as a Monotonic Function of Information for fMRI Nucleus Clusters

We showed in the above paragraphs that, in a Voronoï tessellation of an fMRI image, of particular interest is the presence of *maximal nucleus clusters* (MNC), *i.e*., clusters with nuclei having the highest number of adjacent polygons. In this paragraph, we now introduce a measure of the information that MNCs in fMRI images yield. We demonstrate that MNC reveal regions of the brain with higher levels of cortical information in comparison with non-MNC cortical regions, that uniformly yield less information.

In a series of papers, Rényi (Rènyi, 1961; Rènyi, 1966), introduced a measure of information of a set random events. Let *X* be a set random events such as the occurrence of polygonal areas in a Voronoï tessellation and let *β* > 0, *β* ≠ 1, *p*(*x*) the probability of the occurrence of *x* in *X*. Then Rényi entropy *H*_*β*_(*X*) is defined by

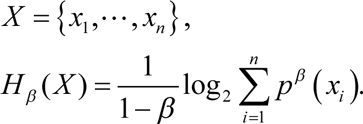

Because of the relationship between Rényi entropy of a set of events and the information represented by events, Rényi entropy and information are interchangeable in practical applications (Rènyi, 1982; Bromiley et al., 2010). In fact, it has been shown that Rényi entropy *H*_*β*_(*X*) is a monotonic function of the information associated with *X*. This means that Rényi entropy can be used as a measure of information for any order *β* > 0(27).

Let *X*_*MNC*_, *X*_*nonMNC*_ be sets of MNC polygon areas and non-MNC polygon areas in a random distribution of tessellation polygon areas. Also, let 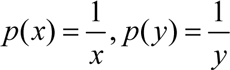 be the probability of occurrence of *y* ∈ *X*_*MNC*_, *y* ∈ *X*_*nonMNC*_. Notice that the nuclei in MNCs have the highest concentration of adjacent polygons, compared all non-MNC polygons. Based on measurements of Rényi entropy for MNC vs. non-MNC observations, we have confirmed that Rényi entropy of nucleus polygon clusters is consistently higher than the set of non-MNC polygons (**Figures 2 E-G**). This finding indicates that MNCs yield higher information than any of the polygon areas outside the MNCs.

In sum, Rényi entropy provides a measure of the information in maximal nucleus clusters and the surrounding zones of fMRI images. This means that the information from areas occupied by MNCs vs. non-MNC areas can be measured and compared. This also means that the maximal nucleus clusters are equipped with higher entropy values (and corresponding higher information), which contrasts with the measure of information in the surrounding non-MNC zones. Hence, MNCs make it possible to pinpoint the highest source of information in fMRI images.

**Figure 1.**
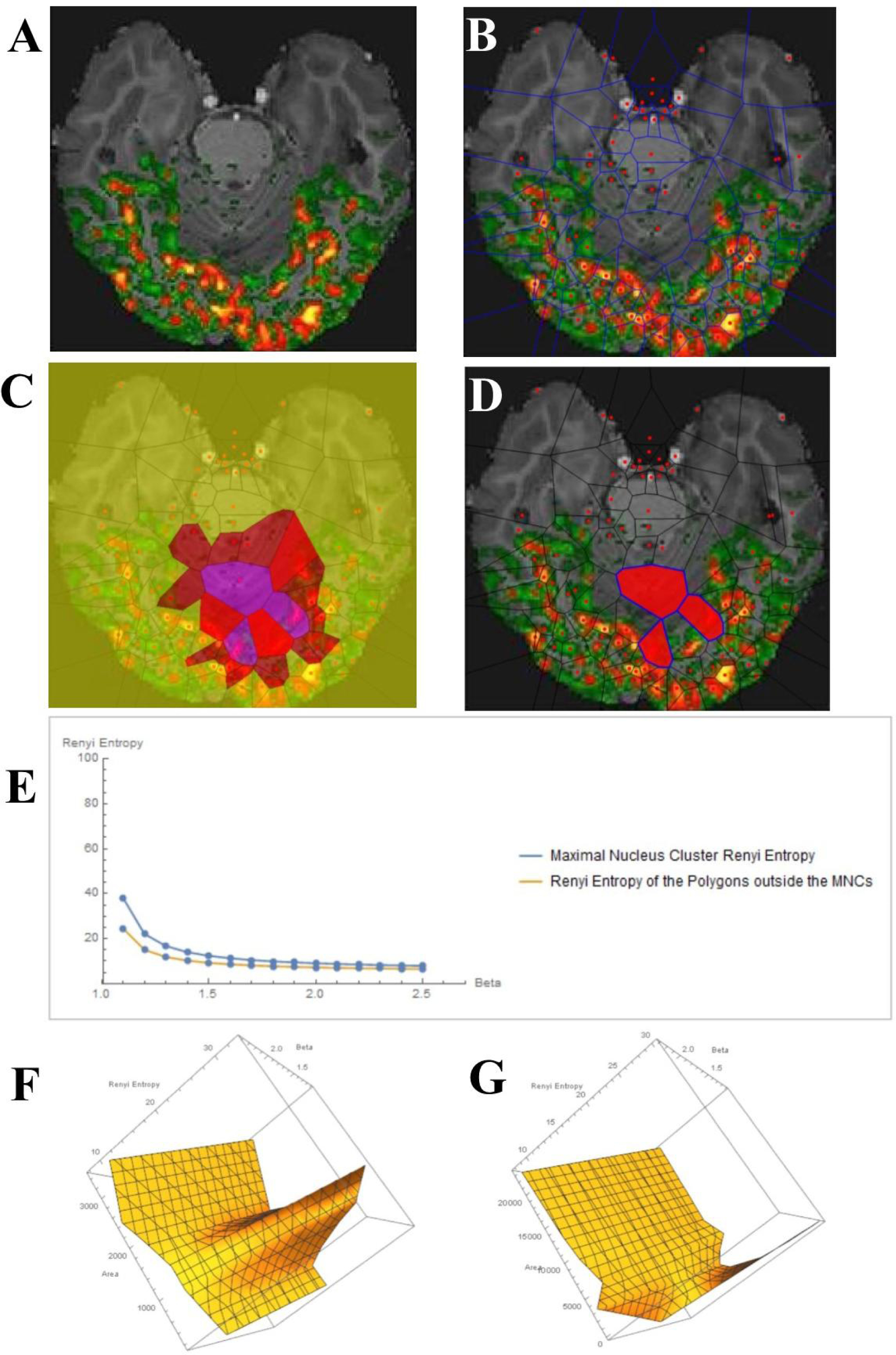
An fMRI image (**Figure 2A**), taken from Mandelkow et al. (2016), underwent Voronoï tessellation (**Figure 2B**). Each •in the tiling represents a generating point with particular features, such gradient orientation and brightness. There are no two • that have the same description. For this reason, every Voronoï region has a slightly different shape. **Figure 2C** displays areas with the higher number of adjacent polygons, in this case three. **Figure 2D** shows a sample maximal nucleus cluster *N* for a particular generating point represented by the dot • in N. In this Voronoï tiling, the nucleus N has several adjacent (strongly near) polygons. Since N has the highest number of adjacent polygons, it is maximal. This *N* is of particular interest, since the generating point • in N has *a gradient orientation that is different from the gradient orientation of any other generating point in this particular tiling*. **Figure 2E** shows Rényi entropy values of maximal nucleus clusters, compared with the surrounding areas of fMRI images. The *x* axis displays the values of the Beta parameter for 1.1 ≤ *β* ≤ 2.5. **Figures 2F and G** display Rényi entropy values vs. number polygon areas vs. 1.1 ≤ *β* ≤ 2.5 of, respectively, MNC and polygons outside the MNC. MNC Nuclei surrounded by polygons with smaller areas have higher Rényi entropy, which tells us that smaller MNC areas yield more cortical information than MNCs with larger areas.

## RESULTS

Each frame from Mandelkow et al. (2016) showed that stimulus-correlated fMRI activity is widespread in the occipital and ventral temporal cortices, consistent with their established involvement in basic vision and object recognition. For each frame, we produced tessellated images with one or more maximal mesh regions. Furthermore, we generated tessellated images showing one or more MNC. The MNC were found to be located in anterior or central brain zones, far from the posterior primary visual sensory areas. The entropy values are higher in the associative zones than in the primary sensory ones. It means that the associative area, during vision, display more activity than the surrounding ones, including the primary visual area. See **Figure 3** for further details.

**Figure 3.**
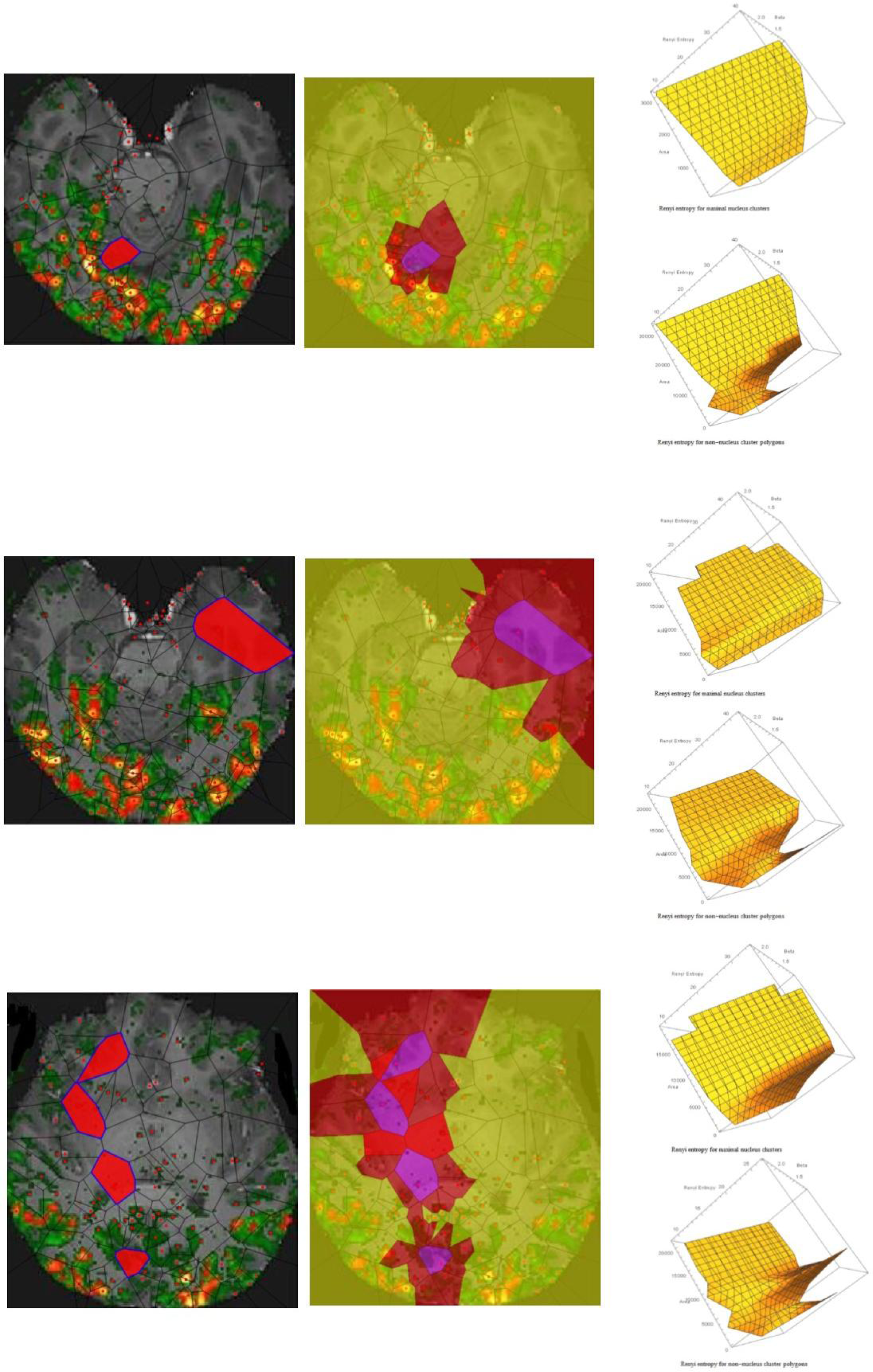
The left side of the Figure depicts MNC tessellations of three samples taken from Mandelkow et al. (2016). The right part displays, for every sample, the Rényi entropy values vs. number polygon areas of MNC (upper part) and outside MNC (lower part)

## DISCUSSION

The current paradigm describes an objective hierarchical landscape of cognition, where a relatively linear and continuous increase in simboli content occurs over network-depth. A continuum of stratified landscapes takes place in a hierarchically ordered connectome, where a graded order of class is represented by a diagram of connected nodes. Starting from external inputs, an emergent aggregation of functions and representations gives rise to hierarchical abstractions processing, where *abstraction* is defined as a process creating general concepts of representation, by emphasizing common features (Taylor et al., 2015). A structural pyramid of the human connectome with gross network-depth directionality can be sketched.*Concrete* regions at the base of pyramid, connected to the outside world, are related to simple perceptions, sensory processing or physical actions.*Abstract* regions at the pinnacle of the pyramid, which are deepest in the brain network and least connected with the outside world, are related to more abstract concepts and symbols. Behavioral elements are conceptualized as brain functions, tasks or behaviors. It means that even abstract behaviors start with sensory inputs and the information representation flow converges towards less complex schemes, e.g., the ideas (Pexman et al. 2016). Unified concepts tend to compress and amalgamate the information content of an idea or an observable event, in order to retain only information which is relevant for an individualized goal or action (Taylor et al., 2015).

Here we proposed and tested an alternative hypothesis, where the above mentioned pyramid of complexity is inverted. The abstract concepts are more intricate and contain more information than the external world. The brain does not *extract* and concentrate the meaning of external objects in simple ideas, rather it *dilutes* the meaning of external objects in complicated ideas. The information is not *aggregated*, rather it is *scattered* in the brain. Recent papers suggest that the brain activity might be multidimensional and shaped in guise of a donut-like structure. It means that the brain function lies in a dimension higher than the environment. Brain activity can be compared to a phase space where projections take place among regions temporarily equipped with different functional dimensions, each one mapping the other. In the powerful topologic framework of BUT and its variants, the world external to the observer *is* a single set of objects, while the world internal to the observer *is* many sets of objects with matching description. Biologically significant, environmental inputs’ single descriptions become matching descriptions in our thoughts. The object’s concept lies in a dimension higher than the object. It means that, e.g., mental images of cats are not pale copies of cats. While mental images are matching descriptions, cats are single descriptions. Thus, contrary to common belief, the object’s concept is more intricate than the object. The brain contains more dimensions and symmetries than the environment and encompasses all the concepts its symmetries might display. We might speculate that such species-specific pool of central nervous system’s symmetries, despite their finite number, can be arranged in countless, versatile modular structures, giving rise to the range of possibilities available for the human brain.

According to the BUT dictates, single descriptions grasp less information and entropy than matching descriptions. Indeed, the energy-BUT variant describes an increase of energy from single descriptions to higher projections. It means that, if our model is true, the primary sensory areas might display less entropy than the associative areas. Therefore, to test our hypothesis, we needed to evaluate the changes in entropy values in fMRI images, in given timescales and in different areas. We used a novel image analysis technique. The major new elements in the evaluation of fMRI images are nucleus clusters, maximal nucleus clusters, strongly near maximal nucleus clusters, convexity structures that occur whenever max nucleus clusters intersect. We showed that in a Voronoï tessellation of an fMRI image, of particular interest is the presence of *maximal nucleus clusters* (MNC), *i.e*., clusters with the highest number of adjacent polygons. We demonstrated that MNC reveal regions of the brain with higher levels of cortical information in comparison with nonMNC cortical regions, that uniformly yield less information. We showed that, in touch with our hypothesis, such brain zones with higher levels of cortical information correspond to associative areas.

In sum, a progressive symmetry reconstruction might occur in human brain from the lower-dimensional single features of the *concrete* periphery to the higher dimensional matching features of the *abstract* center. It means that brain concepts and ideas are from *above*, not from *below*.

